# Genome-wide association study reveals sex-specific genetic architecture of facial attractiveness

**DOI:** 10.1101/339226

**Authors:** Bowen Hu, Ning Shen, James J. Li, Hyunseung Kang, Jinkuk Hong, Jason Fletcher, Jan Greenberg, Marsha R. Mailick, Qiongshi Lu

**Affiliations:** Department of Statistics, University of Wisconsin-Madison, Madison, WI, USA 53706; Department of Psychology, University of Wisconsin-Madison, Madison, WI 53706; Waisman Center, University of Wisconsin-Madison, Madison, WI, USA 53705; La Follette School of Public Affairs, University of Wisconsin–Madison, Madison, WI 53706; Department of Sociology, University of Wisconsin–Madison, Madison, WI 53706; School of Social Work, University of Wisconsin–Madison, Madison, WI 53706; Department of Biostatistics and Medical Informatics, University of Wisconsin-Madison, Madison, WI, USA 53792

**Keywords:** GWAS, facial attractiveness, Wisconsin Longitudinal Study

## Abstract

Facial attractiveness is a complex human trait of great interest in both academia and industry. Literature on sociological and phenotypic factors associated with facial attractiveness is rich, but its genetic basis is poorly understood. In this paper, we conducted a genome-wide association study to discover genetic variants associated with facial attractiveness using 3,928 samples in the Wisconsin Longitudinal Study. We identified two genome-wide significant loci and highlighted a handful of candidate genes, many of which are specifically expressed in human tissues involved in reproduction and hormone synthesis. Additionally, facial attractiveness showed strong and negative genetic correlations with BMI in females and with blood lipids in males. Our analysis also suggested sex-specific selection pressure on variants associated with lower male attractiveness. These results revealed sex-specific genetic architecture of facial attractiveness and provided fundamental new insights into its genetic basis.

## Introduction

Facial attractiveness is a complex human trait of great interest in sociology, psychology, and related fields due to its profound influence on human behavior. Although variability exists across individuals and cultures, it has been suggested that some commonly agreed cues are used by people everywhere to judge facial beauty [1, 2]. As a trait that is well integrated into people’s daily life experience, it is unsurprising that facial attractiveness influences a variety of sociological outcomes. Studies have suggested that facial attractiveness is associated with job-related outcomes [3–6], academic performance [7], and economic mobility [8]. It affects human psychological adaptations and serves as an important influence of mate preference and reproductive success [9–14]. Even attractive babies receive more nurturing from their mothers than unattractive babies [15]. Further, people all over the world highly prize beauty. The annual revenue of the cosmetic industry is around 18 billion dollars in the US [16]. Fashion and beauty dominate daily discussions on traditional media as well as social media posts. Understanding the perception of attractiveness is a great interest in both academia and industry.

The genetic basis of facial attractiveness may provide new and mechanistic insights into this complex human trait. Evidence suggested that attractiveness is heritable and genetic variations explain a substantial fraction of its variability [17, 18]. However, no genetic variant or gene underlying the biology of facial attractiveness has been identified to date. Our understanding of its genetic architecture is certainly far from complete. Well-phenotyped facial attractiveness data from the Wisconsin Longitudinal Study (WLS) have expanded our knowledge on the complex relationship between facial attractiveness and various sociological traits including educational aspirations and occupation [19–24]. Recently, dense genotype data have been made available in WLS. These advances make it possible for the first time to identify specific genetic components associated with facial attractiveness and probe its genetic architecture.

We performed a genome-wide association study (GWAS) on 3,928 samples from WLS to identify single-nucleotide polymorphisms (SNPs) associated with facial attractiveness. In addition, sex-stratified analyses suggested distinct genetic architecture between the perception of male and female attractiveness. Integrated analysis of GWAS results and transcriptomic and epigenetic functional annotations also provided mechanistic insights into how genetics may influence facial attractiveness.

## Results

### GWAS identifies genetic loci associated with facial attractiveness

We conducted a GWAS for facial attractiveness on individuals of self-reported European ancestry in WLS. After quality control, a total of 3,928 individuals and 7,251,583 SNPs were included in the analyses (**Supplementary Table 1**). In 2004 and 2008, each individual’s high-school yearbook photo was independently rated by 12 coders (6 females and 6 males) who were selected from the same birth cohort as the WLS participants. These scores were normalized into two metrics to represent the average attractiveness ratings from female and male coders on each individual (**Methods**). We conducted two separate GWAS on all samples using attractiveness scores given by female and male coders as quantitative traits (**Figure 1A** and **Supplementary Figures 1-2**). These two analyses are referred to as FC-AS (female coders, all samples) and MC-AS (male coders, all samples) throughout the paper. We identified one genome-wide significant locus at 10q11.22 for FC-AS (rs2999422, p=9.2e-10; **Table 1** and **Figure 2A**). No genome-wide significant loci were detected for MC-AS. Additionally, we identified three loci showing suggestively significant associations (p<1.0e-6) with FC-AS (6p25.1) and MC-AS (20q13.11 and 2q22.1; **Supplementary Figures 3-4**).

**Figure 1.**
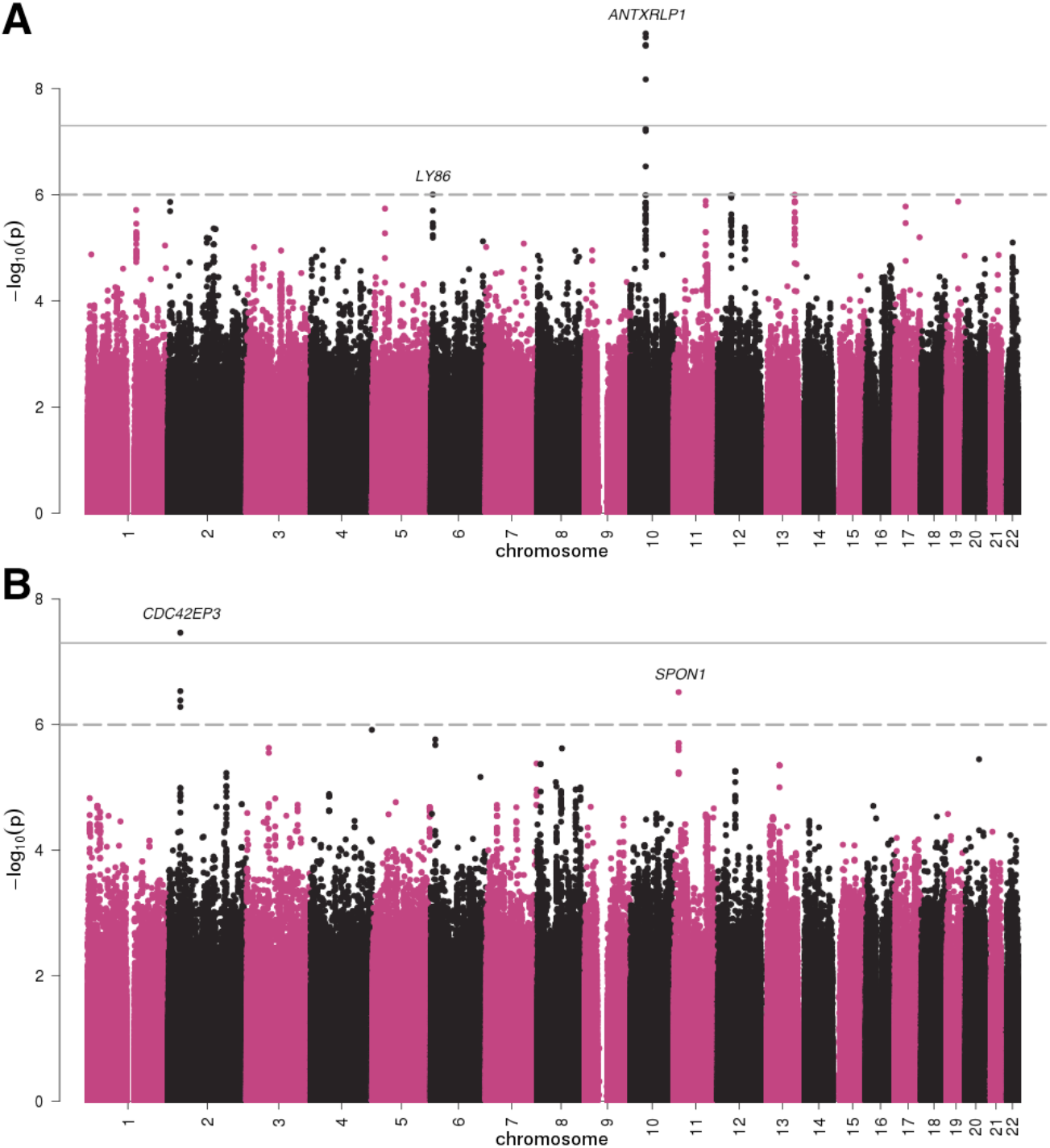
Manhattan plots for (A) FC-AS and (B) MC-FS analyses. The horizontal lines denote the genome-wide significance cutoff of 5.0e-8 and a suggestive cutoff of 1.0e-6.

**Figure 2.**
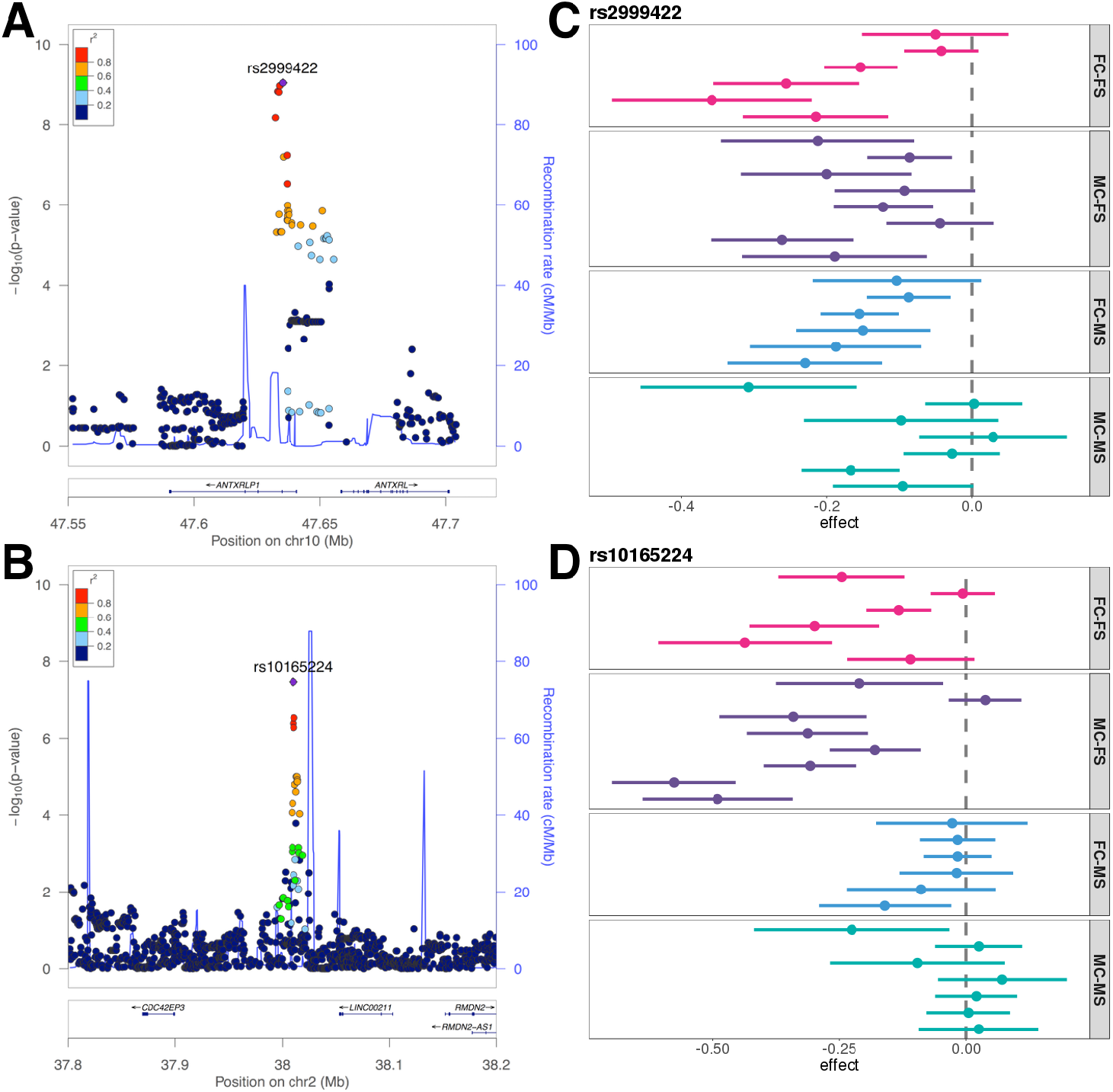
Associations at two genome-wide significant loci. (**A**) Association with FC-AS at 10q 11.22. (**B**) Association with MC-FS at 2p22.2. (C-D) Association signals for leading SNPs at these two loci across coders. Each interval shows association with attractiveness score given by one coder. Error bars denote standard error of effect estimates. Only coders who rated more than 500 male or female samples were included in the analysis.

**Table 1:**
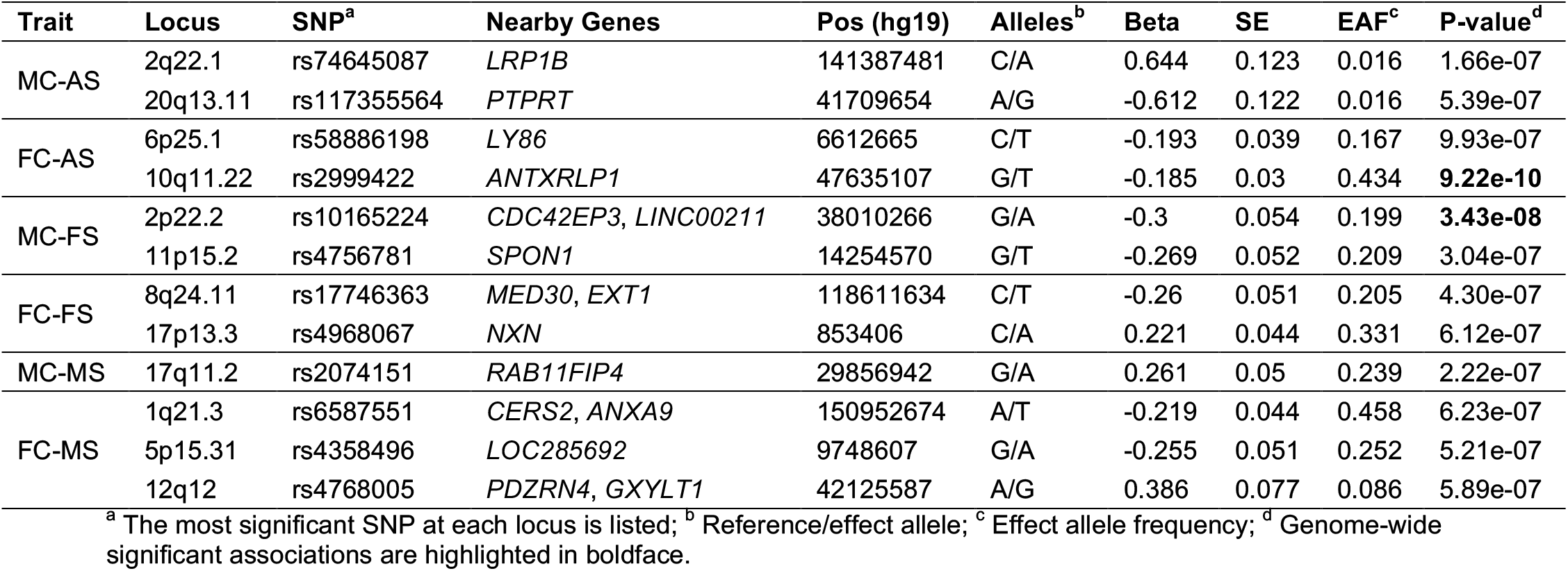
Genetic loci associated with facial attractiveness.

Attractiveness is known to have sex-specific associations with various social factors [25–27]. Thus, we hypothesized that different genetic components may be associated with male and female attractiveness and the genetic architecture may further diverge when comparing the perception from males and females. We conducted four sex-specific GWAS (**Figure 1B** and **Supplementary Figures 5-6**) based on female coders and female samples (FC-FS), female coders and male samples (FC-MS), male coders and female samples (MC-FS), and male coders and male samples (MC-MS). We identified one additional genome-wide significant locus at 2p22.2 for MC-FS (rs10165224, p=3.4e-8; **Figure 2B**). Seven loci at 1q21.3, 5p15.31, 8q24.11, 11p15.2, 12q12, 17p13.3, and 17q11.2 showed suggestive associations in sex-stratified analyses (**Table 1** and **Supplementary Figures 7-10**). Of note, associated loci identified for FC-AS and MC-AS all showed consistent effects in sex-specific analyses (**Supplementary Table 2**).

A total of 80 coders participated in the attractiveness study in WLS (**Methods**). To investigate heterogeneity of identified signals, we performed association analyses using attractiveness scores from each coder separately. In order to maintain statistical power, we focused on coders who rated more than 500 male or female samples in WLS (**Supplementary Figure 11**). The genome-wide significant association identified for FC-AS, i.e. rs2999422, showed consistently negative associations with attractiveness scores from all female coders and most male coders (**Figure 2C**). The genome-wide significant locus in MC-FS analysis, rs10165224, also showed consistency – negative associations were identified for all male coders except one (**Figure 2D**). Consistent association patterns were also observed for other identified loci (**Supplementary Figure 12**). These results suggest that associations identified in our analyses were not driven by several coders’ particularly strong opinions. Rather, they represent genetic associations with the consensus of opinions among coders.

### Heritability and selection signatures of facial attractiveness

Consistent with many other complex traits [28], top associations identified in our analyses only explained a small fraction of phenotypic variability. We obtained positive estimates of chip heritability for FC-AS (0.109) and MC-aS (0.277) using genome-wide data [29]. However, we note that standard errors for these estimates were high (0.149 for FC-AS and 0.159 for MC-AS) and the GREML algorithm [29] did not converge in sex-specific analyses, likely due to limited sample size. Next, we applied linkage disequilibrium (LD) score regression [30] to partition heritability by tissue and cell type. Interestingly, several tissues related to reproduction and hormone production were strongly enriched for heritability of facial attractiveness (**Supplementary Table 3**). Despite not reaching statistical significance after correcting for multiple testing, testis was the top tissue for FC-AS (enrichment=3.9, p=0.04) and ovary was the most enriched tissue for MC-AS (enrichment=4.5, p=0.032), MC-FS (enrichment=5.7, p=0.040), and FC-FS (enrichment=3.0, p=0.005). Reproductive organs were not highlighted in male-specific analyses (i.e. MC-MS and FC-MS).

Further, we investigated the relationship between minor allele frequencies (MAF) and minor allele effects on facial attractiveness. We grouped SNPs with MAF between 0.05 and 0.5 into 10 equally-sized bins based on MAF quantiles. Interestingly, minor alleles with low frequencies tend to have negative effects on male facial attractiveness (**Figure 3**). The mean minor allele effect on FC-MS from SNPs in the lowest 10% MAF quantile was −0.005, implying very strong statistical evidence for its deviation from zero (p=7.3e-313; two-sided t-test). SNPs in the highest 10% MAF quantile, however, did not show significantly negative associations (mean=-4.1e-5, p=0.493). This hinted at selection pressure on genetic variants associated with negative male attractiveness. The selection signature in females was not as clear.

**Figure 3.**
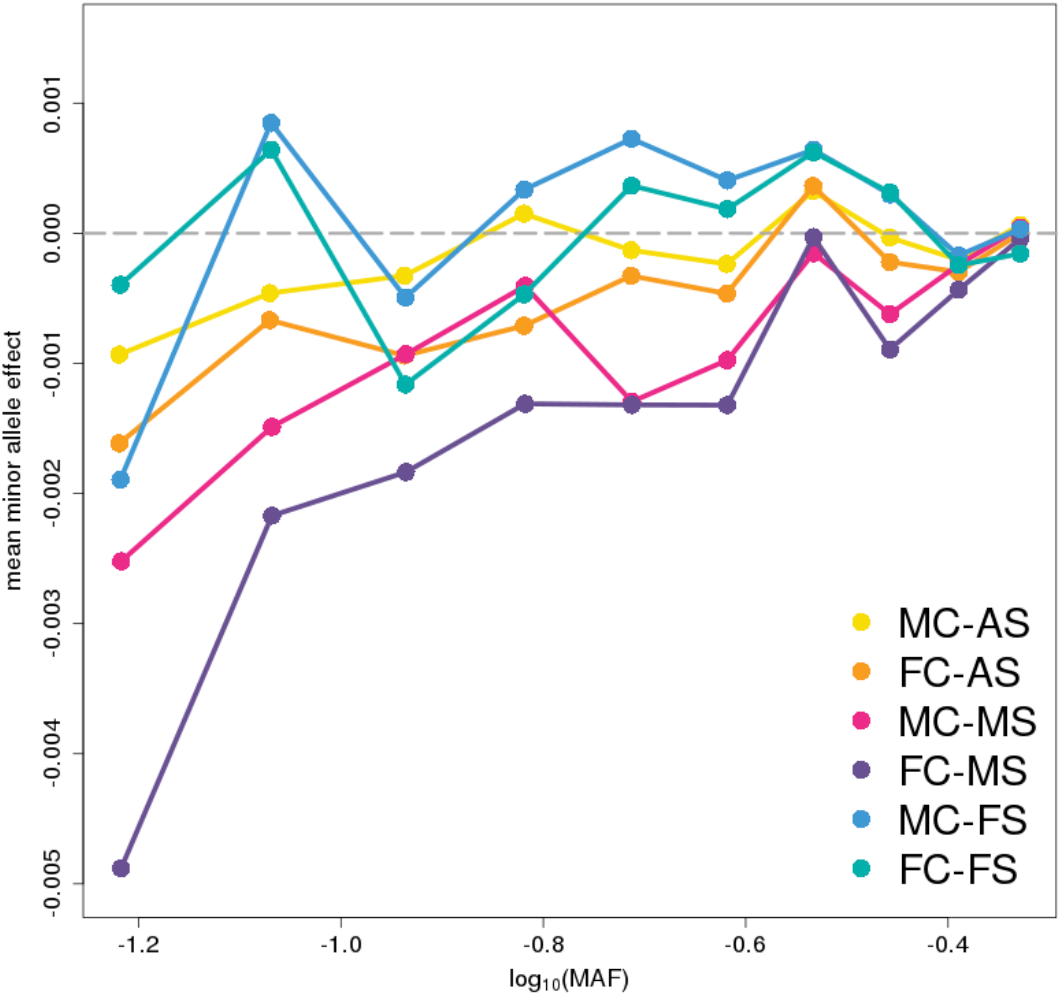
Selection signatures of facial attractiveness. SNPs with MAF between 0.05 and 0.5 were grouped into 10 equally-sized bins based on MAF quantiles. For SNPs in each bin, average MAF was calculated shown on the x-axis, while the average minor allele effects are shown on the y-ax¡s.

### Candidate genes at identified loci

One genome-wide significant locus at 10q11.22 was identified for FC-AS. The leading SNP at this locus, rs2999422, is located in an intron of pseudogene *ANTXRLP1*. The closest protein-coding gene is *ANTXRL* (**Figure 2A**). Interestingly, both *ANTXRL* and *ANTXRLP1* have substantially higher expression levels in testis than in other tissues (**Supplementary Figure 13**). In females, *ANTXRL* is highly expressed in fallopian tube, breast, and cervix while *ANTXRLP1* is not specifically expressed in reproductive organs. This locus has been previously reported to associate with skin pigmentation [31] and transferrin saturation [32]. The genes closest to the three suggestively significant loci for FC-AS and MC-AS, i.e. *LRP1B, PTPRT*, and *LY86* (**Supplementary Figures 3-4**), have been reported in multiple association studies. Specifically, *LRP1B* is a member of the low-density lipoprotein (LDL) receptor family and is associated with body-mass index (BMI) [33], aging [34], and age at menarche [35]; *PTPRT* is associated with facial morphology [36]; *LY86* is associated with waist-hip ratio [37] and hip circumference [38].

Among the loci identified in sex-stratified analyses, one locus at 2p22.2 reached genome-wide significance for MC-FS (**Figure 2B**). The leading SNP rs10165224 is located in an intergenic region between protein-coding gene *CDC42EP3* and RNA gene *LINC00211*. This locus was known to be associated with height [37, 39]. Genes at the seven suggestively significant loci (**Supplementary Figures 7-10**) were also associated with various traits related to facial features. Both *SPON1* at 11p15.2 and *NXN* at 17p13.3 are associated with facial morphology [36]. *NXN* is also associated with vulvitis and vulva disease (MCIDs: VLV008 and VLV036). Additionally, *EXT1* at 8q24.11 is associated with obesity [40] and *PDZRN4* at 12q12 is associated with BMI [41] and skin pigmentation [31]. The locus at 1 q21.3 contains a large LD block covering multiple genes, among which *ANXA9* is highly expressed in skin tissue (**Supplementary Figure 14**) and is associated with melanoma [42]. Finally, *TAS2R1* at 5p15.31 is specifically expressed in testis tissue in males (**Supplementary Figure 15**).

A few leading SNPs at identified loci are expression quantitative trait loci (eQTL) for nearby genes (**Supplementary Table 4**). The genome-wide significant SNP at 10q11.22 for FC-AS, rs2999422, is an eQTL for *ANTXRL* across various tissues (minimum p=1.1e-8); rs17746363 is an eQTL for *MED30* in skeletal muscle (p=2.5e-6); rs2074151 and rs6587551 are eQTL in thyroid for *RAB11FIP4* (p=1.2e-5) and *CTSS* (p=5.6e-26), respectively. To systematically utilize multi-tissue eQTL data and better quantify associations at the gene level, we performed cross-tissue transcriptome-wide association analyses for six facial attractiveness traits using the UTMOST method (**Methods; Supplementary Figure 16**) [43]. We identified four significant gene-level associations after correcting for multiple testing: *SYT15* at 10q11.22 for FC-AS (p=1.0e-6), *CTSS* at 1q21.3 for FC-MS (p=9.6e-7), *RPL22* at 1p36.31 for MC-FS (p=1.5e-7), and *ATAD5* at 17q11.2 for MC-MS (p=8.9e-7). *SYT15* is 700kb upstream of *ANTXRL*, the genome-wide significant locus for FC-AS. *CTSS* is located at a suggestively significant locus for FC-MS and is associated with BMI [44]. *ATAD5* is 700kb upstream of the suggestively significant locus for MC-MS and is known to associated with many complex traits including height, waist circumference, hip circumference, and BMI [45–47]. *RPL22* is a novel association and is highly expressed in multiple reproductive organs including ovary, uterus, and cervix (**Supplementary Figure 17**). All gene-level associations for six attractiveness traits are summarized in **Supplementary Table 5**.

### Relationship between facial attractiveness and other complex traits

Next, we investigated the relationship between facial attractiveness and various related human traits. First, we compared our results with association signals for several human facial features including facial morphology [48], skin pigmentation [49], and hair color [50]. Although top SNPs identified in these studies did not show overall significant enrichment for associations with facial attractiveness, some SNPs showed consistent effects in our analyses (**Supplementary Tables 6-8**). A leading SNP associated with facial morphology, rs1523446, showed consistently negative associations with all six attractiveness traits (e.g. p=6.8e-4 for MC-AS) and is known to associate with waist-hip ratio across populations [45, 51]. SNP rs35096708, a leading association locus for skin pigmentation, vitiligo, and hair color [52, 53] also showed consistent associations in all our analyses (e.g. p=0.002 for MC-AS and 0.006 for FC-AS). A recent study identified more than 100 associated loci for hair color [50], many of which also showed associations with attractiveness in our analyses. For example, a SNP associated with lighter hair color – rs34764931, was positively associated with all six attractiveness traits (e.g. p=0.006 for FC-AS).

Analyses based on comparing top associations across multiple traits have limited statistical power given moderate sample size and do not capture the polygenic architecture. To further reveal the relationship between facial attractiveness and other complex traits, we estimated genetic covariance between facial attractiveness and 50 complex traits with publicly accessible GWAS summary statistics which covered a spectrum of social, psychiatric, anthropometric, metabolic, and reproductive phenotypes (**Supplementary Table 9**). Results for all 300 pairs of genetic covariance are summarized in **Supplementary Table 10**. One pair of traits – female BMI (BMI-F) and MC-FS, showed strong and negative correlation, and the p-value achieved Bonferroni-corrected significance (covariance=-0.053, p=4.7e-5). Additionally, three other pairs of traits showed genetic covariance with Benjamini-Hochberg false discovery rate (fdr) below 0.1, including BMI and MC-FS (covariance=-0.035; p=6.4e-4), high-density lipoprotein cholesterol (HDL-C) and FC-MS (covariance= −0.058; p=8.2e-4), and total cholesterol (TC) and FC-MS (covariance=-0.063; p=5.1e-4). Interestingly, female attractiveness traits were negatively correlated with all three BMI traits, and the correlation signal was the strongest when attractiveness was rated by male coders (i.e. MC-FS) and the BMI analysis was specific to females (**Figure 4A**). However, such a relationship was completely absent between male attractiveness and bMi. In fact, both FC-MS and MC-MS were positively correlated with BMI traits although the p-values were non-significant. In contrast, genetic covariance between attractiveness and lipid traits was specific to male samples, especially the FC-MS analysis in which female coders’ scores were analyzed (**Figure 4B**).

**Figure 4.**
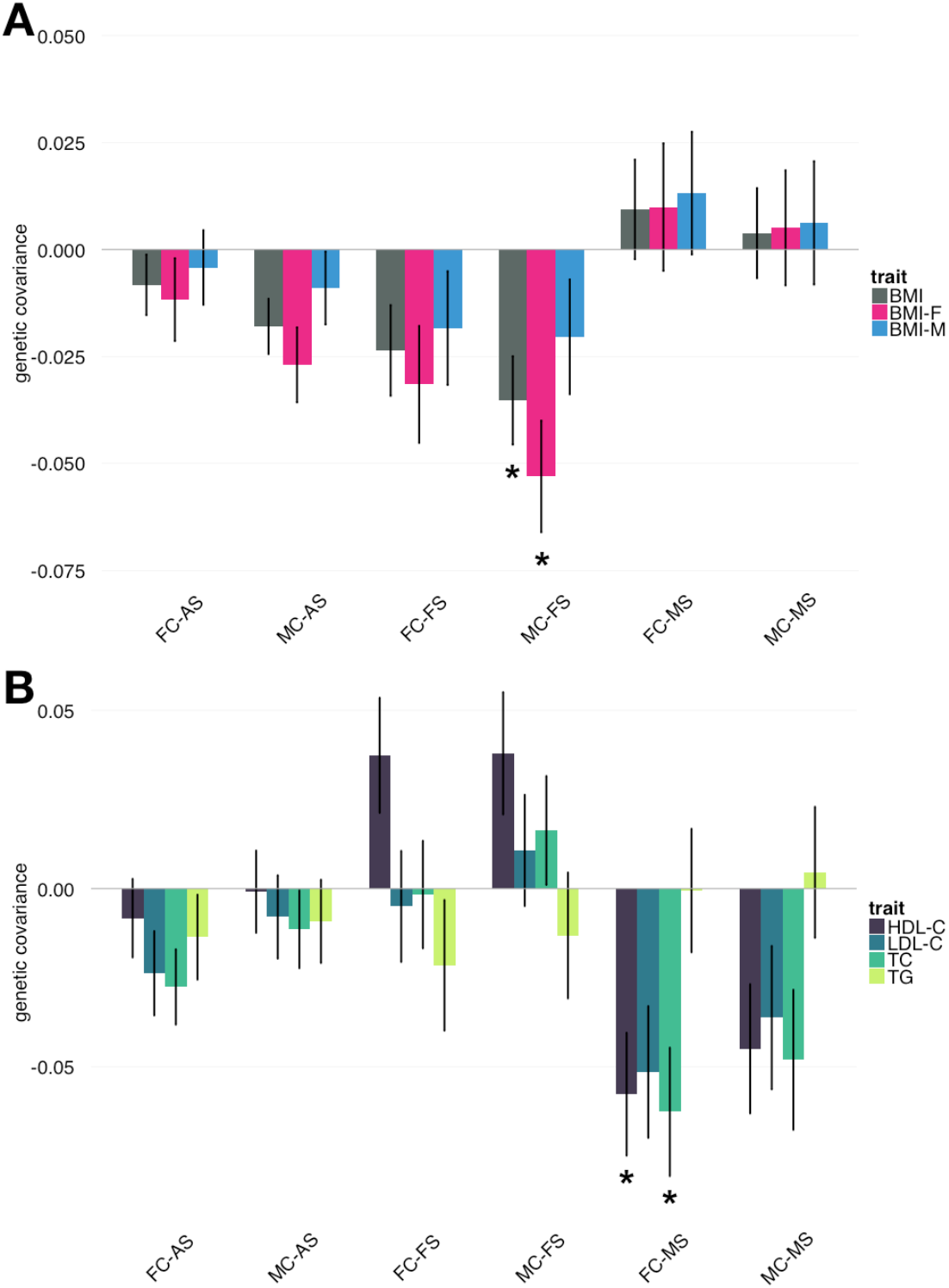
Genetic covariance between facial attractiveness and (A) BMI traits, (B) lipid traits. Intervals show the standard error of covariance estimates. Genetic covariance with fdr < 0.1 are marked by asterisks.

Furthermore, we explored if the strong genetic covariance of MC-FS with BMI-F could be explained by a causal relationship between these traits. We performed robust Mendelian randomization (MR) to infer causality (**Methods**). We identified a negative effect from BMI-F to MC-FS with p=0.099. Our results hinted at a causal effect between BMI-F and MC-FS but the validity of the relationship requires future investigation using larger samples.

## Discussion

Despite tremendous interests from both academia and industry, the genetic basis of facial attractiveness is poorly understood, partly due to the scarcity of well-phenotyped facial attractiveness in large-scale cohorts with genetic information. Carefully-measured human facial attractiveness, in conjunction with dense genotype data in WLS, made it possible to identify specific genetic components for facial attractiveness. In this paper, we conducted a GWAS to identify DNA variants associated with human facial attractiveness. We identified two genome-wide significant loci on 10q11.22 and 2p22.2 and highlighted several genes via eQTL analysis and transcriptome-wide association analysis. Multiple candidate genes identified in our analyses were specifically expressed in human tissues involved in reproduction and hormone production, and this pattern was further confirmed in genome-wide heritability enrichment analysis. Additionally, loci strongly associated with facial attractiveness were enriched for known associations with BMI and waist-hip ratio. Through comparing leading associations across multiple studies, we also identified potentially pleiotropic loci shared with facial morphology, skin pigmentation, and hair color. Further, via a genome-wide genetic covariance estimation approach, we identified strong evidence for shared genetic architecture of facial attractiveness with BMI and lipid traits. Of note, sex-specific genetic architecture of facial attractiveness was a recurrent pattern observed in almost all our analyses. The loci that reached significance in analyses based on all samples showed consistent effects between males and females, but sex-specific analyses revealed a list of new loci. In multi-trait analyses, only female attractiveness (especially MC-FS) showed strong and negative genetic correlation with BMI while male attractiveness was more strongly correlated with blood cholesterol levels which are known to be involved in the synthesis of testosterone and other steroid hormones [54]. Finally, variants and genes were identified for both male and female attractiveness but the selection pressure on negative associations of male attractiveness seemed to be particularly strong. These results not only provided fundamental new insights into the genetic basis of facial attractiveness, but also revealed the complex relationship between attractiveness and a variety of human traits.

This study was not without limitations. First, although WLS provided a great opportunity to study the genetics of facial attractiveness, the sample size was moderate and we could not find an independent cohort to replicate our association findings due to the uniqueness of this phenotype. Due to weak effect sizes, extreme multiple testing, and ubiquitous confounding, replication and validation are critical steps in studies of complex trait genetics. We have performed a variety of analyses to assess the heterogeneity of identified associations, including comparing association signals between males and females as well as across different coders. Still, validity of our findings remains to be further investigated using independent samples. Second, attractiveness measurements in WLS were based on high-school yearbook photos. Each photo was graded by 12 different coders, thereby creating robust average measurements. However, we note that this phenotyping approach does not cover every aspect of attractiveness. What are the roles of age, physical body shape, and facial expression in the perception of attractiveness? And what is the shared and distinct genetics between attractiveness and closely related facial phenotypes such as symmetry, averageness, and sexually dimorphic features [14]? These are just a handful of questions beyond the scope of this study. Additionally, we note that both the raters and the people being rated were from one state that was racially and ethnically quite homogeneous and we only included samples with European ancestry in this study. Further, the yearbook photos in WLS were collected more than sixty years ago. It is unclear how well our results can be generalized to other populations, age groups, and generations. If the same high school yearbook photos were to be rated for facial attractiveness by a more ethnically or racially diverse set of raters, and if the findings were to be replicated, then the inference regarding genetic association of attractiveness would be strengthened. Nevertheless, this study was a successful attempt to pin down genetic components of human facial attractiveness. Many of these unanswered questions will be exciting directions to explore in the future. We have little doubt that robust and comprehensive phenotypic measurements, coupled with larger sample sizes from diverse populations, will further advance our understanding of this interesting human trait.

## Methods

### WLS data details

WLS is a longitudinal study of a 1/3 random sample of over ten thousand Wisconsin high school graduates in 1957. Facial attractiveness in WLS was measured based on each individual’s 1957 high school year book photo by 12 coders (six females and six males) selected from the same cohort in 2004 and 2008. An 11-point rating scale was used to quantify attractiveness. End-points of rating were labeled as “not at all attractive” and “extremely attractive” for 1 and 11, respectively. In this paper, we used normalized average ratings from female coders and normalized average ratings from male coders as two quantitative traits for facial attractiveness. Normalization was performed in a prevailing fashion as subtracting mean and then divided by standard deviation.

Genetic data were obtained from saliva samples in the year of 2006 and 2007 using Oragene kits and a mail-back protocol. We used genotype data imputed against the Haplotype Reference Consortium (HRC) panel. All participants provided informed consent under a protocol approved by the Institutional Review Board of the University of Wisconsin-Madison. Individuals were removed if they did not satisfy one of the following criteria: 1) genotype missingness rate was less than 0.05 in all chromosomes, 2) surveyed sex matched with genetic sex, 3) surveyed relatedness matched with genetic relatedness 4) the individual was not an outlier in respect of heterozygosity or homozygosity, 5) the individual was not an ancestral outlier, 6) attractiveness ratings were available, 7) the individual had self-reported European ancestry and 8) born between 1937 −1940. SNPs were removed if: 1) it had a call rate less than 0.95, 2) Hardy-Weinberg exact test p-value was less than 1.0e-5, 3) minor allele frequency was below 0.01, or 4) imputation quality score was below 0.8. After quality control, 7,251,583 autosomal SNPs and 3,928 individuals remained in the data.

### GWAS analysis

In the analysis using all samples (i.e. MC-AS and FC-AS), we applied linear mixed model (LMM) implemented in the GCTA software [55] to perform association analysis while correcting for relatedness among samples. In addition, sex, round of coding (i.e. was attractiveness rated in 2004 or 2008), dummy variables for birth year were included in the model as covariates. In sex-stratified association analyses, we applied linear regression instead of LMM due to the reduction in sample size and the consequent non-convergence of the restricted maximum likelihood algorithm and add the first two principal components into covariates. WLS data were collected on high school graduates of the same year and distant cousins may be involved due to the study design. To adjust for family structure in linear regression analysis, we kept only one individual in each pair of samples with relatedness coefficient greater than 0.05. Relatedness coefficients among samples were estimated using PLINK [56]. After these additional quality control steps, 1, 792 male samples and 2,062 female samples remained in sex-stratified analyses, PLINK was used to perform association analysis with sex, round of coding, birth year, and the first two principal components included as covariates.

### eQTL data and transcriptome-wide association analysis

Multi-tissue gene expression and eQTL data were acquired from data portal of the Genotype-Tissue Expression (GTEx) project (https://www.gtexportal.org). We applied UTMOST [43] to perform cross-tissue transcriptome-wide association analysis for six facial attractiveness traits. We used cross-tissue gene expression imputation models trained from 44 tissues in GTEx [57]. Gene-level association meta-analysis was performed using generalized Berk-Jones test [58] implemented in UTMOST software. Statistical significance was determined using a Bonferroni corrected p-value cutoff 3.2e-6.

### Heritability and multi-trait analysis

The GREML method implemented in GCTA software was used to estimate heritability of facial attractiveness [59, 60]. We applied stratified LD score regression [61] implemented in the LDSC software to perform heritability enrichment analysis and identify biologically relevant tissues for facial attractiveness. Tissues with sample sizes greater than 100 in GTEx were included in the analyses. In sex-stratified analyses, non-existent tissues (e.g. testis for females and ovary for males) were removed from the analyses. For each tissue, functional annotation was defined as regions near highly expressed genes (within 50,000 bp up- or downstream). We used median transcripts per million (TPM) as the criterion to select top 10% highly expressed genes. We then estimated partitioned heritability using functional annotation for each tissue while including 53 baseline annotations in the model. P-values were calculated using z-scores of regression coefficient as previously suggested [61].

Leading associations for facial morphology, skin pigmentation, and hair color were acquired from related publications [48–50, 52]. All genome-wide significant loci identified in the discovery stage for facial morphology were included in our analysis. Meta-analysis p-values and p-values from Rotterdam study were used to prioritize SNPs for hair color and skin pigmentation, respectively. Only the most significant SNP at each locus was included in the analysis. A locus was removed if the leading SNP was not present in our study. We used the GNOVA method [62] to estimate genetic covariance between complex traits. Association statistics of six facial attractiveness traits were jointly analyzed with publicly accessible GWAS summary statistics for 50 complex traits (**Supplementary Table 9**). Since samples in WLS were not used in those 50 published datasets, uncorrected genetic covariance estimates were reported in our analyses. Additionally, due to numerically unstable estimates for heritability, we report genetic covariance instead of genetic correlation throughout the paper.

We used MR-Egger [63] approach implemented in ‘MendelianRadomization’ R package to perform causal inference between complex traits. We selected instrumental SNP variables by applying a LD cutoff of 0.05 and a p-value cutoff of 1.0e-9. Based on these criteria, 31 top SNPs for BMI were included in our analysis.

### Other bioinformatics tools

Manhattan plots and QQ plots were generated using ‘qqman’ package in R [64]. Locus plots for GWAS loci were generated using LocusZoom [65].

### Data availability

Genotype data from WLS are available to the research community through the dbGaP controlled-access repository at accession phs001157.v1.p1. Phenotypic data in WLS can be accessed via the WLS data portal https://www.ssc.wisc.edu/wlsresearch/. Summary statistics for six facial attractiveness traits will be publicly released upon publication.

## Acknowledgements

This study was supported by the Clinical and Translational Science Award (CTSA) program, through the NIH National Center for Advancing Translational Sciences (NCATS), grant UL1TR000427. We thank Pamela Herd, Geoffrey Hayes, Joe Savard, Carol Roan, and Kamil Sicinski for their assistance in WLS data curation. We also thank Maryam Zekavat for her assistance in interpreting the involvement of cholesterols in steroid hormone synthesis.

## Competing financial interests

The authors declare no competing financial interests.

## Author contribution

Q.L. designed the study.

B.H. and Q.L. accessed, cleaned, and processed genetic data B.H., N.S., and Q.L. performed statistical analyses.

J.L., J.H., J.G., and M.M. assisted data collection and processing.

H. K. advised on causal inference.

J.F., J.G., and M.M. advised on sociological issues.

Q.L. advised on statistical and genetic issues.

B.H. and Q.L. wrote the manuscript.

All authors contributed in manuscript editing and approved the manuscript.

